# Restoring Function After Severe Spinal Cord Injury Through Bioluminescence-Driven Optogenetics

**DOI:** 10.1101/710194

**Authors:** Eric D. Petersen, Erik D. Sharkey, Akash Pal, Lateef O. Shafau, Jessica R. Zenchak, Alex J. Peña, Anu Aggarwal, Mansi Prakash, Ute Hochgeschwender

## Abstract

The ability to manipulate specific neuronal populations of the spinal cord following spinal cord injury (SCI) could prove highly beneficial for rehabilitation in patients through maintaining and strengthening still existing neuronal connections and/or facilitating the formation of new connections. A non-invasive and highly specific approach to neuronal stimulation is bioluminescent-optogenetics (BL-OG), where genetically expressed light emitting luciferases are tethered to light sensitive channelrhodopsins (luminopsins, LMO); neurons are activated by the addition of the luciferase substrate coelenterazine (CTZ). This approach utilizes ion channels for current conduction while activating the channels through application of a small chemical compound, thus allowing non-invasive stimulation and recruitment of all targeted neurons. Rats were transduced in the lumbar spinal cord with AAV2/9 to express the excitatory LMO3 under control of a pan-neuronal or motor neuron-specific promoter. A day after contusion injury of the thoracic spine, rats received either CTZ or vehicle every other day for 2 weeks. Activation of either interneuron or motor neuron populations below the level of injury significantly improved locomotor recovery lasting beyond the time of stimulation. Utilizing histological and gene expression methods we identified neuronal plasticity as a likely mechanism underlying the functional recovery. These findings provide a foundation for a rational approach to spinal cord injury rehabilitation, thereby advancing approaches for functional recovery after SCI.

## Introduction

The manipulation of specific neuronal populations of the spinal cord following spinal cord injury (SCI) could prove highly beneficial for rehabilitation in patients. This could work by maintaining and strengthening existing neuronal connections and/or facilitating neuronal growth and the formation of new synapses in a controlled, activity dependent manner. Stimulation of circuits in the spinal cord would ideally be highly cell type specific and non-invasive. Electrical stimulation presents a straight-forward means to activate neurons of the spinal cord and although showing clinical promise, this approach has several critical limitations. Electrical stimulation excites all cells within the electrode vicinity, potentially diluting or negating the effect of targeted stimulation of specific beneficial cell types. Electrical stimulation also results in rapid muscle fatigue by preferentially recruiting large, rapidly adapting motor units, limiting the on-time for stimulation*(1),(2)*. Further, electrical stimulation requires a chronic implant, potentially increasing risk to the patient.

Optogenetics is a promising pre-clinical method for stimulating neurons of the spinal cord and overcomes some of the problems with electrical stimulation, allowing activation of specific channels or effectors that can be targeted to discrete, genetically unique neural sub-populations. While used extensively to interrogate neuronal circuits of the brain approaches to enable manipulation of neurons and other cell types in the periphery have only recently been developed *(5)*. However, the need for invasive chronic optical fiber implants connected to an external light source or implanted LED modules poses problems for long-term treatment when applied to the spinal cord. Furthermore, light from an external source is limited to efficiently penetrating the tissue ∼200 µm below the dorsal surface of the spinal cord, thus not reaching the ideal target neurons for stimulation of central pattern generators and motor circuits *(3, 4)*. Another concern for long term optogenetic stimulation as a therapeutic approach is the side effect of high intensity light exposure to neuronal tissue. Heat produced by the illumination source can rapidly raise local temperatures in neuronal tissue by several degrees with commonly used light intensities, affecting neuronal activity and inflammatory activation of astrocytes and microglia *(6, 7)*.

BioLuminescent-OptoGenetics (BL-OG) is a recently developed approach that has the potential to overcome the barriers to clinical success presented by traditional optogenetic approaches for rehabilitation following SCI. BL-OG uses powerful optogenetic elements that do not require an external implant, but instead use light generated internally by tethering bioluminescent luciferases to light sensitive channelrhodopsins, luminopsins (LMO). The bioluminescent light is produced by the breakdown of a specific enzymatic substrate, in this case coelenterazine (CTZ). Stimulation only occurs when the CTZ is injected, producing bioluminescent light through catalysis by the luciferase, resulting in the activation of the opsin. This approach takes advantage of both opto- and chemogenetic concepts by utilizing ion channels for current conduction while activating the channels through the application of a chemical compound, thus allowing non-invasive stimulation and recruitment of all targeted actuators as opposed to only those that can be reached by light from a physical source *(8–15)*. Here we use LMO3 which consists of slow burn *Gaussia* luciferase fused to *Volvox* channelrhodopsin 1. This system has been demonstrated to consistently activate expressing neuronal cells with little to no off target effects caused by the substrate and bioluminescent reaction that takes place *(16, 17, MGR)*. Moreover, bioluminescence is light emitted without heat (“cold light”) and thus does not approach the damaging levels encountered for traditional optogenetics *(18)*. Utilizing LMOs for neural stimulation in the spinal cord presents an innovative approach for activating neurons that might be therapeutically beneficial to recovery following SCI that was not previously possible with other approaches.

Here we sought to determine if genetically targeted stimulation restricted to neurons of the lumbar spinal cord or specifically to motor neurons of the lumbar spinal cord would be beneficial for locomotor recovery following experimental spinal cord injury.

## Results

### Bioluminescent optogenetic stimulation of spinal neurons

To assess the possibility of BL-OG stimulation re-engaging neurons below the site of a spinal cord injury we transduced neurons of the lumbar enlargement with AAV vectors to express LMO3 (Fig. 1A). At the time of AAV injection, we also implanted a lateral ventricle cannula for easy application of the luciferase substrate CTZ (Fig. S1). Contusion injury of the thoracic spinal cord was carried out 3 weeks later, followed by BL-OG stimulation and testing of locomotor behavior (Fig. 1B). LMO3 expression was under control of a pan-neuronal human synapsin promoter (hSyn) or a motoneuron-specific rat Homeobox 9 promoter (Hb9). Expression under the hSyn promoter was consistently concentrated to neurons located within laminae 4-8 and 10, with some expression in lamina 9 (Fig. 1C). The Hb9 promoter successfully restricted expression almost exclusively to motor neurons in lamina 9 (Fig. 1D), with some interneurons also expressing the construct, which is consistent with previous reports *(19–21)*. In the rostral – caudal dimension LMO3 expression with both promoters was observed throughout the majority of the lumbar enlargement, with the highest levels of expression closest to the injection site.

**Fig. 1.**
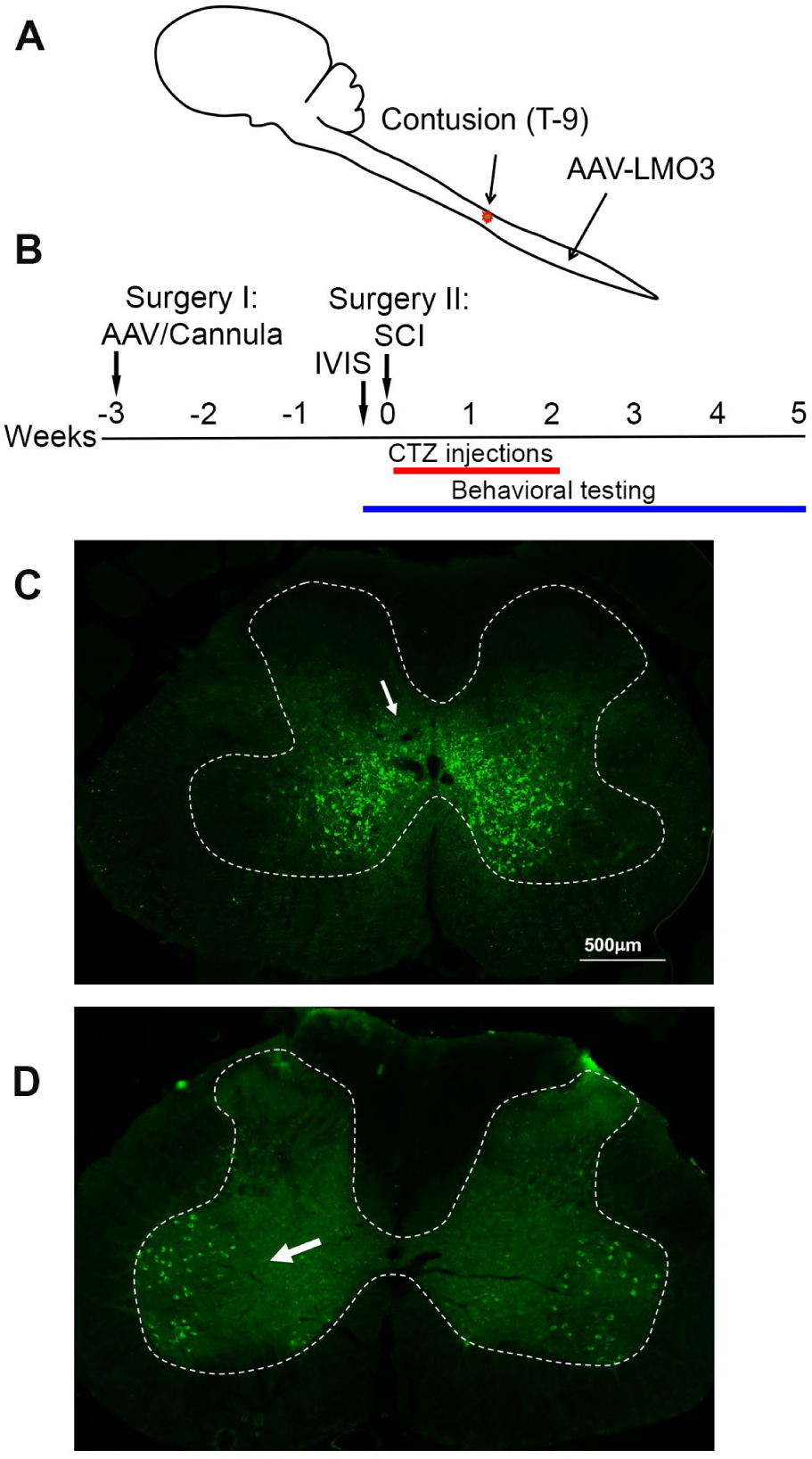
Spinal cord injury model. (**A**) Schematic of the experimental model with viral injection for BL-OG stimulation in the lumbar enlargement and contusion injury in the thoracic region. (**B**) Timeline of experimental procedures with the first surgery for lateral ventricle cannula placement and virus injection three weeks prior to injury. (**C**) Expression of AAV 2/9 hSyn-LMO3 in the lumbar spinal cord (arrow pointing to expressing interneurons). The highest levels of expression are restricted to interneuron populations in lamina 4-8 and 10 with some expression more dorsal and in lamina 9. (**D**) Expression of AAV 2/9 Hb9-LMO3 in the lumbar spinal cord (arrow pointing to expressing motor neurons). The Hb9 promoter restricts expression to motor neurons in lamina 9. Some low level of expression does occur throughout other lamina of the cord.

To insure that viral transductions resulted in LMO3 expression at sufficient levels and in the intended anatomical region, we took advantage of the unique feature of LMOs allowing for *in vivo* bioluminescence imaging. Bioluminescence was detected over the lumbar region of the spinal cord in rats transduced with LMO3 (Fig. 2A). Light intensities over time consistently peaked between 10 and 30 minutes post CTZ application and decayed over the next hour (Fig. 2B). Utilizing *in vivo* bioluminescent imaging not only allowed us to confirm LMO3 expression, we were also able to verify proper cannula function. As we performed *in vivo* bioluminescent imaging prior to the contusion injury, we avoided continuing with animals that had insufficient expression or an improper working cannula.

**Fig. 2.**
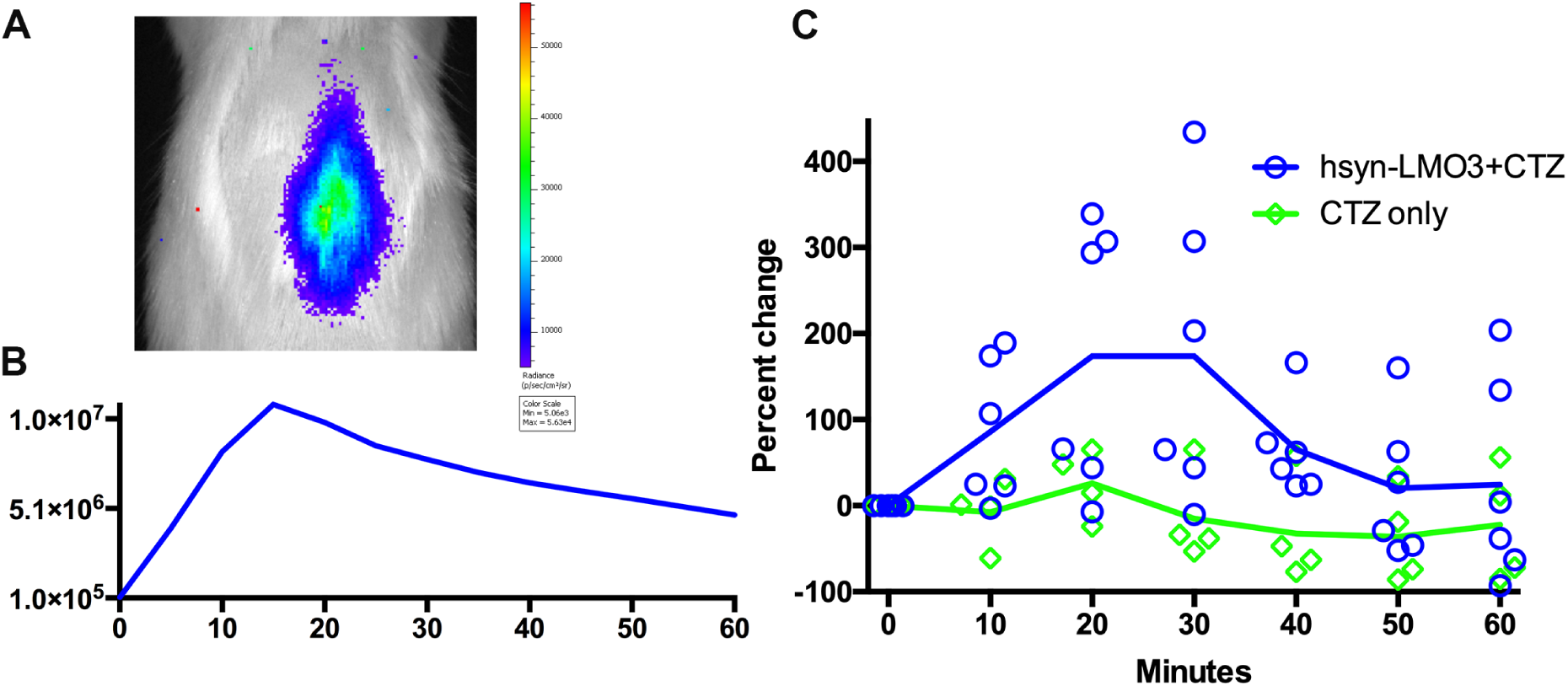
Bioluminescent optogenetic stimulation of spinal cord neurons. (**A**) Example of *in vivo* bioluminescent imaging of a rat expressing LMO3 in the lumbar spinal cord following CTZ infusion through the lateral ventricle. Luminescence is localized over the lumbar region of the cord. (**B**) A representative trace of luminescence over time following CTZ infusion. (**C**) Single unit electrophysiological response in the lumbar spinal cord of rats expressing AAV 2/9 hSyn-LMO3 compared to non-expressing animals when CTZ is infused through the lateral ventricle. Similar to luminescence over time, activity increases and peaks between 10 and 30 minutes following CTZ infusion. n= 7 for LMO3 expressing and n= 4 for non-expressing animals.

Next we determined the effect of LMO stimulation on activity of spinal neurons when CTZ is infused through the ventricle. We confirmed that increases in neuronal activity within the lumbar spinal cord followed a similar timeline as observed with *in vivo* bioluminescent imaging (Fig 2C). When testing the electrophysiological effect of CTZ on naïve rats under the same recording conditions, we did not find any change in activity over baseline spiking rates.

### Bioluminescent optogenetics results in accelerated and enhanced locomotor recovery after SCI

All animals expressing LMO3 in the lumbar spinal cord had Basso, Beattie, and Bresnahan (BBB) ratings of 21 (perfect gait) before undergoing surgery for spinal cord injury. After thoracic contusion injury rats were randomly assigned to two groups with one group receiving CTZ and the other group receiving vehicle via ventricular infusion. Applications were delivered every other day for 14 days, starting the day after SCI surgery. SCI rats that received CTZ mediated neural stimulation showed a significant improvement in locomotor scores which persisted even after the treatment period (Fig. 3A). Animals that received stimulation via hSyn-LMO3 (n=6) had a final mean BBB score of 13.2, representing animals with frequent to consistent weight supported plantar steps and frequent front limb-hindlimb coordination. Animals that received stimulation via Hb9-LMO3 (n=6) had a mean BBB score of 11.4, representing animals able to take frequent to consistent weight supported steps. Those treated with the vehicle solvent (n=11) had a final mean BBB score of 7.9, representing animals that are able to move both hindlimbs in a sweeping motion without any weight support. Animals receiving BL-OG mediated stimulation regardless of the neuronal population targeted improved at a faster rate than vehicle treated controls, with significantly better locomotor scores from days 7-35 post injury. For locomotor recovery scores, there was a significant main effect for treatment (F(2,20)=18.148, p=2.22×10^−5^) and a significant main effect for time (F(2.839,154)=228.33, p=3.89×10^−31^). There was also a significant interaction effect for treatment by time point (F(5.68,154)=6.400, p=4.61×10^−5^). At the experimental endpoint, 100% of rats that received stimulation were able to take weight bearing steps (BBB of 10 or higher) while only 18% of vehicle treated animals were able to take weight bearing steps (Fig. 3B). We also found animals that received stimulation tended to regain bladder control sooner than the vehicle treated group, however this difference was not significant (Fig. S2).

**Fig. 3.**
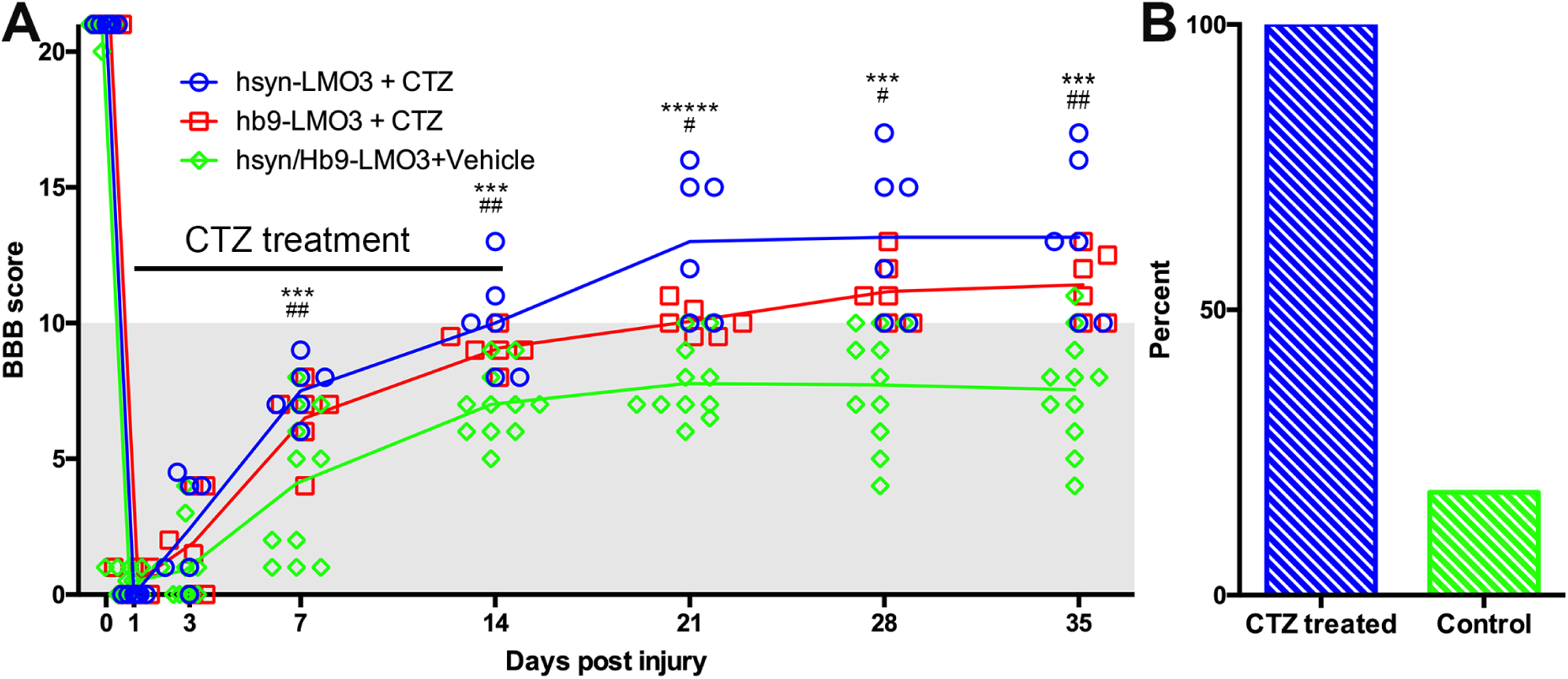
Accelerated and enhanced locomotor recovery with BL-OG. (**A**) BBB locomotor scores following injury and treatment for animals that received neuronal stimulation (hSyn-LMO3+CTZ), motor neuron stimulation (Hb9-LMO3+CTZ) or vehicle following injury (no stimulation). Those which received stimulation regardless of neuronal subpopulation recovered at a faster rate, to a greater extent, and maintained their status following the treatment period compared to the vehicle treated group. For Bonferroni post hoc: *= p<0.05, **=p<0.01, ***=p<0.001, ****=p<0.0001, *****=p<0.00001, for hSyn-LMO3+CTZ vs vehicle and #= p<0.05, ##=p<0.01, ###=p<0.001 for Hb9-LMO3+CTZ vs vehicle. n= 6 for hSyn-LMO3 + CTZ, n=6 for Hb9-LMO3 + CTZ, n= 11 for vehicle treated animals. (**B**) Comparison of the percentage of weight supporting animals at the endpoint for those that received BL-OG stimulation compared to vehicle.

In an independent experiment, we sought to determine if locomotor recovery could be explained by off target effects of bioluminescence. For this, we tested whether bioluminescence produced by the luciferase sbGluc and substrate CTZ without an optogenetic channel present could impact recovery. We used the construct hSyn-sbGLuc-B7-EYFP where B7 is a transmembrane domain replacing the optogenetic channel so the luciferase is extracellular and tethered to the cell membrane as in the LMO constructs. This tested the potential effects of bioluminescence, CTZ, and all breakdown products from the chemical reaction. Following the same injury and treatment protocol used for LMO expressing animals, we found no difference at any time point between sbGLuc-B7-EYFP expressing animals receiving CTZ and vehicle (Fig. S3). Both treatment groups recovered in a similar manner as the vehicle treated LMO3 group.

### Bioluminescent optogenetics effects recovery after SCI through increasing neuronal plasticity

The positive effect of post-injury engagement of neurons could be based on different mechanisms or a combination thereof. Since the treatment used was an early intervention, it could have influenced the extent of degeneration at the lesion site in a variety of ways. To assess sparing of myelinated white matter at the site of injury, eriochrome cyanin staining was used. We found no differences between either stimulation treatment condition and the vehicle treatment in the cross sectional area of preserved white matter at the lesion epicenter (Fig. 4).

**Fig. 4.**
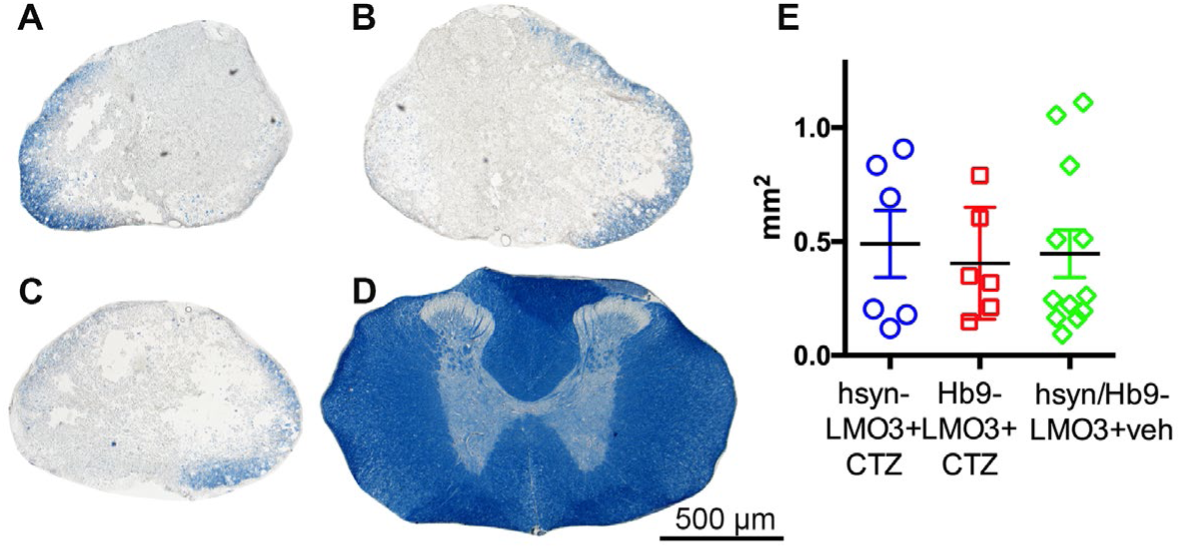
No sparing of white matter at the injury site. (**A-D**) Cross sections of spinal cords stained with eriochrome cyanin which stains white matter blue. A: hSyn-LMO3+CTZ; B: Hb9-LMO3+ CTZ; C: Vehicle treated; D: Example of the white matter present in the same region of the spinal cord in a non-injured rat. (**E**) Comparison of the cross sectional area of spared white matter following injury and treatment. There were no differences in the amount of degeneration that occurred as a result of the contusion injury with or without neural stimulation.

Based on these results, we performed experiments to determine if BL-OG stimulation is able to influence neuronal plasticity under the conditions used in this study. For this, we measured mRNA levels in the region stimulated (lumbar) at eight days post injury. The experiment was carried out as described above with fresh tissue collection on day 8 post injury, when rats had received four CTZ applications. When assessing gene expression at this time point we found animals that received stimulation tended to have higher levels of markers for neuronal plasticity: GAP-43, MAP2, PSD-95, and NMDAR2d (Table 1). We also compared expression levels for genes associated with inflammation and apoptosis to determine if either of these have a role in promoting recovery. We did not find consistent trends for either inflammation or apoptotic markers when comparing animals treated with CTZ versus vehicle. To further assess the effect of BL-OG stimulation on gene expression and towards developing a rational approach to non-invasive stimulation for neurological disorders, we assessed relative transcript levels with RNAseq in the lumbar and thoracic regions of the spinal cord. We found that overall, there was a strong correlation of genes expressed in the thoracic region, the site of injury, between the treated and vehicle group (Fig. S4, Table S2). There was a much weaker correlation of genes expressed in the lumbar region, the site of stimulation, between the treatment and vehicle groups (Fig. S5, Table S3). A variety of genes that potentially have roles in supporting neuronal activity and health were upregulated in the lumbar region of the treatment group. We also identified genes in both the lumbar and thoracic regions that represent an improved inflammatory state for the treatment group (Fig. S6 and S7).

**Table 1.** Gene expression levels after injury and treatment. Comparison of relative mRNA levels for various biomarkers in the lumber spinal cord following injury and subsequent treatment. All levels are expressed as fold change over non injured sham animals. N=5 for all groups except for MAP2, PSD-95,VEGF, iNOS, Caspase 3, and BCL-1, where one animal from the sham group did not have detectable levels of the gene of interest and was excluded. For Arginase 1, one animal from sham and one from vehicle did not have a detectable level of the gene of interest and was excluded.

| Marker | Function | Fold of sham (SE) | ANOVA | Bonferroni post hoc |
| --- | --- | --- | --- | --- |
| GAP-43 | Axon growth | CTZ: 6.49 (1.49)<br>Veh: 2.99 (1.65)<br>Sham: 1.00 | p=0.034<br>F=4.523,14 | CTZ - veh: p=0.248<br>CTZ - sham: p=0.035<br>Veh - sham: p=0.909 |
| MAP2 | Dendrite growth | CTZ: 12.56 (3.49)<br>Veh: 3.65 (1.55)<br>Sham: 1.00 | p=0.014<br>F=6.405,13 | CTZ - veh: p=0.059<br>CTZ - sham: p=0.020<br>Veh - sham: p=1.000 |
| PSD-95 | Post synaptic structure | CTZ: 5.07 (1.43)<br>Veh: 1.53 (0.77)<br>Sham: 1.00 | p=0.037<br>F=4.520,13 | CTZ - veh: p=0.093<br>CTZ - sham: p=0.065<br>Veh - sham: p=1.000 |
| NMDAR2d | Post synaptic receptor | CTZ: 6.85 (2.30)<br>Veh: 1.79 (0.75)<br>Sham: 1.00 | p=0.023<br>F=5.271,14 | CTZ - veh: p=0.071<br>CTZ - sham: p=0.031<br>Veh - sham: p=1.000 |
| BDNF | Neurotrophic factor | CTZ: 0.56 (0.14)<br>Veh: 0.75 (0.09)<br>Sham: 1.00 | p=0.001<br>F=13.352,14 | CTZ - veh: p=0.122<br>CTZ - sham: p=0.001<br>Veh - sham: p=0.043 |
| VEGF | Neurotrophic factor | CTZ: 17.33 (4.33)<br>Veh: 3.55 (1.90)<br>Sham: 1.00 | p=0.029<br>F=4.964,13 | CTZ - veh: p=0.156<br>CTZ - sham: p=0.034<br>Veh - sham: p=1.000 |
| iNOS | M1 microglia/macrophage | CTZ: 8.98 (1.22)<br>Veh: 5.70 (2.45)<br>Sham: 1.00 | p=0.028<br>F=5.005,13 | CTZ - veh: p=0.586<br>CTZ - sham: p=0.027<br>Veh - sham: p=0.269 |
| Arginase1 | M2 microglia/macrophage | CTZ: 11.48 (5.62)<br>Veh: 7.24 (6.37)<br>Sham: 1.00 | p=0.118<br>F=2.662,12 | CTZ - veh: p=0.225<br>CTZ - sham: p=0.246<br>Veh - sham: p=1.000 |
| Caspase 3 | Apoptotic | CTZ: 1.31 (0.29)<br>Veh: 1.26 (0.89)<br>Sham: 1.00 | p=0.487<br>F=0.768,13 | CTZ - veh: p=0.950<br>CTZ - sham: p=0.914<br>Veh - sham: p=1.000 |
| Bcl-2 | Anti-apoptotic | CTZ: 1.85 (0.48)<br>Veh: 0.60 (0.36)<br>Sham: 1.00 | p=0.264<br>F=1.507,13 | CTZ - veh: p=0.349<br>CTZ - sham: p=.0889<br>Veh - sham: p=1.000 |

## Discussion

The majority of spinal cord injuries are contusion injuries that leave behind areas of intact neural tissue below the site of injury. Neurons from those areas often maintain intact connections, even across the site of injury, yet patients are paralyzed. To explore if these functionally dormant populations could be re-engaged in participating in active spinal circuitry resulting in improved functional output, we stimulated neurons below the site of a contusion injury for two weeks using a bioluminescent optogenetics approach. We found that stimulating neurons with this approach resulted in greatly improving the rate and extent of locomotor recovery following injury.

Optogenetic stimulation following spinal cord injury has been tested in a mouse model of cervical SCI previously and was found capable of improving breathing following treatment *(22)*. However, the practical challenges of light delivery to often centrally located spinal cord target populations in a moving organ has hampered application of this approach to SCI. In principle, chemogenetic approaches are ideal for manipulation of spinal cord neurons as they require application of a small chemical compound to activate genetically targeted neurons without requirements for hardware. In one study, Chen et al. restored stepping ability in mice with staggered bilateral hemisections by administering a KCC2 agonist. The same result was achieved by selective expression of hyperpolarizing DREADDs (hM4Di) in inhibitory interneurons between and around the staggered spinal lesions *(23)*. However, chemogenetic approaches to more clinically relevant SCI injury models are lacking. Using a rat spinal cord contusion model we explored BL-OG, a chemogenetic approach that takes advantage of using ion channels for current conduction rather than GPCRs, thus making it independent of requirements for expression of specific GPCR coupled pathways.

Using BL-OG, we found that stimulating either primarily interneurons or primarily motor neurons resulted in significant functional improvements at faster rates and to a greater extent in CTZ treated, stimulated animals compared to vehicle treated control animals. Of note is that for the motor neuron stimulation paradigm (Hb9-LMO3), relatively few neurons expressed the LMO construct compared to the pan neuronal paradigm (hSyn-LMO3), yet a very similar end result was achieved. Our findings of improvement regardless of the neuronal population that was stimulated is different from those in another study where directly reducing the excitability of inhibitory Vgat interneurons, but not directly increasing the excitability of Vglut excitatory interneurons resulted in improvement after bilateral hemisections *(23)*. Putting the differences in SCI model and species aside, the results from the two studies are not incompatible. Rather, they point to a general theme potentially underlying recovery after SCI, that of pushing the ratio of excitation to inhibition towards excitation. In our study we favored excitation by stimulating motor neurons or globally interneurons, while the other study did so by tempering the excessive activity of inhibitory interneurons, with both approaches re-engaging functionally dormant neuronal pathways to contributing to movement after an SCI. It will be of great interest to determine if genetically targeting stimulation with BL-OG to other genetically distinct neural subpopulations could further improve outcomes following SCI. For example, there exist a variety of genetically identifiable interneuron subtypes that make up mammalian central pattern generators (CPGs) and which remain to be fully characterized that will likely prove useful targets to promote recovery following SCI *(24)*.

The hypothesis that BL-OG stimulation of locomotor networks can induce remodeling at the neuronal and synaptic level is supported by our results. We did not find an effect on sparing of white matter that could have had a large impact on behavioral outcomes. This led us to directly test the effect of LMO stimulation on markers for plasticity. In RT-PCR experiments we found all four neuronal plasticity markers encompassing axon, dendrite, and synaptic remodeling to be expressed at higher levels after injury but even more so as a result of treatment. We also tested a variety of other biomarkers to determine if inflammation was affected as a result of stimulation and did not find strong evidence to support this. RNA sequencing studies further confirmed lack of effects of lumbar stimulation on injury pathology by showing highly similar gene expression profiles in the thoracic region between vehicle and treatment groups, while gene expression profiles were altered in the lumbar region. All together, we believe that BL-OG mediated recovery following SCI is largely mediated by optogenetically induced neuronal plasticity and potentially maintenance of neural networks and improved inflammatory state. In the previous study of optogenetic stimulation of cervical SCI beneficial effects were largely mediated by neuronal plasticity *(22)*. Consistent with these findings, optogenetics has previously been demonstrated to induce neuronal plasticity *in vitro, in vivo*, and to promote recovery following different types of neural trauma*(22, 25–27)*. Most recently, BL-OG stimulation has been used to restore function following stroke, where the benefits of stimulation were found to be a result of optogenetically induced neuronal plasticity *(17)*.

Our results are highly encouraging in the context of clinical translatability. Viral vectors for gene delivery are increasingly finding their way into the clinic. Coelenterazine has been used without detriment in animal imaging studies for decades. The specific route of application in our study, lateral ventricle infusion, was chosen based on considerations of practicability in rats. For human application, alternate routes would apply (intravenous, intranasal). However, we report on our initial, limited study that needs to be followed up to address several critical issues. For example, although our contusion injury is severe, we don’t know if we reached the limit of effectiveness with our stimulation treatment. Going in the other direction, in cases of less severe contusions and expected higher numbers of preserved intact neurons below the site of injury, it is possible that the pan-neuronal stimulation (hSyn-LMO3) used in this study will show detrimental effects due to simultaneous activation of counteracting CPG populations. Furthermore, it will be important to try the BL-OG stimulation approach at later time points after the occurrence of the injury. Lastly, we might see synergistic effects when combining BL-OG stimulation with physical exercises.

From a translational perspective we expect that our results can be built upon in the future to develop improved approaches to treating SCI that leverage the capacity of optogenetic stimulation for induction of plasticity for successful treatment of human patients. Given the observation in our study that animals that received stimulation tended to regain bladder function sooner, we believe this approach may present a means to improve bladder control if the stimulation was purposely targeted to the nuclei of the cord that are responsible for bladder control. This would be a major quality of life improvement for patients with SCI.

## Materials and Methods

### Animals

Adult female Sprague Dawley rats, 4-6 months of age, bred on site, weighing 280-350 grams were used. All experimental procedures were performed in accordance with guidelines from the NIH and were approved by the Central Michigan University Institutional Animal Care and Use Committee (IACUC). Animals were kept in 12 hour light/dark cycle rooms and fed ad libitum.

### Plasmids and virus

LMO3, the third generation of excitatory LMOs, was expressed in neurons of the lumbar spinal cord utilizing an adeno-associated virus serotype 2/9. LMO3 consists of slow burn *Gaussia* luciferase fused to *Volvox* channelrhodopsin 1, with a yellow fluorescent protein tag and was expressed under the human synapsin (hSyn) promoter or under the rat Homobox 9 (Hb9) promoter. A rat version of the Hb9 promoter described in references *(19–21)*, which is 99% similar to the mouse version was synthesized by Genscript and cloned into the pAAV-hSyn-LMO3 plasmid to replace the hSyn promoter, creating pAAV-Hb9-LMO3 using standard restriction cloning techniques. The B7 transmembrane sequence from the mouse CD80 antigen was cloned into the AAV vector to replace the optogenetic channel *(28)*. Plasmids were confirmed by sequencing. High titer stocks of hSyn-LMO3 virus were made by ViroVek. The other two viruses were made in-house using previously described methods for triple plasmid transfection in HEK293FT cells to encapsulate the constructs in a pseudotyped 2/9 capsid *(12)*.

### Surgery

All surgeries were conducted under aseptic conditions.

#### Lateral ventricle cannulation

The lateral ventricle cannula consists of an infusion cannula (3280PM/SPC cut 4 mm below pedestal, Plastics One) to access the ventricle that is externalized through a PinPort (VABR1B/22, Instech Labs) that allows repeated aseptic access. The two parts are connected by 2.0 cm of 22G polyurethane tubing (VAHBPU-T22, Instech labs) (Fig. S1). For placement, an incision was made to expose the skull, periosteum removed, and bone dried thoroughly. A burr hole was drilled at −1.0 mm from bregma, 1.5 mm right of the midline for insertion of the cannula *(29)*. Three machine screws (00-96 c 3/32, Plastics One) were inserted into hand drilled holes (D69, Plastics One) 0.742mm forward, behind, and to the left of where the infusion cannula would be placed. The infusion cannula was lowered 4 mm below the skull and secured to the skull and screws by dental acrylic. The port was externalized though the skin on the neck and sutured tightly around the base with a 4-0 silk suture, and incisions closed with staples. Cannulas were kept clear from obstruction by infusing saline twice a week prior to the second surgery.

#### Viral injections

During the same surgery, animals received viral injections in the lumbar spinal cord. The spinal cord was exposed by making an incision over the T-13/L1 vertebra and the soft tissue between the two vertebra was cleared to expose a minimal amount of the cord. The spinal column was stabilized using vertebral clamps. The virus was infused using a 10 µL World Precision Instruments syringe with 35G beveled needle. The virus was injected 0.5 mm lateral to the midline and 1.5 mm ventral to the surface at the following volumes per side: 2.5µL at 1×10^13^ copies/mL for hSyn-LMO3, 6 µL at 5×10^12^ copies/mL for Hb9-LMO3, and 2.5 µL at 3 ×10^12^ for hsyn-sbGluc-B7-EYFP; volumes were adjusted to result in equal levels of expression judged by expression of the EYFP reporter. All were infused at a rate of 0.16 µL/minute and left in place for an additional 5 minutes.

#### Spinal cord injury

The spinal cord was exposed with a laminectomy at T-9 and stabilized with vertebral clips. An NYU impactor was aligned with the exposed spinal cord and weight dropped from 25 cm to induce a severe contusion *(30)*. Following surgeries, the incision site over the cord was closed in layers, animals were given 5 mL of lactated Ringers solution, and placed on a heating pad to recover thermoregulation.

### IVIS imaging

Bioluminescence imaging was done under isoflurane anesthesia, with an IVIS Lumina LT (Perkin Elmer) where the CTZ was infused through the cannula and the animal was imaged for a time series with the exposure set at 5 minutes, f-stop at 1, with large binning.

### *In vivo* electrophysiology recordings

Acute recordings were performed under 1.2-1.5g/kg urethane. Animals were secured in a Kopf spinal stereotax and a Hamilton syringe with a 25G beveled needle, loaded with CTZ was lowered into the lateral ventricle. A laminectomy was performed at the L1 vertebra and a 32 channel electrode array (A2×16, NeuroNexus) was lowered on one side of the cord to a depth of 2 mm. For acquisition, a Blackrock Microsystems CerePlex µ head stage and CerePlex Direct acquisition system were used. Recordings were filtered with a 250 Hz high pass fourth order Butterworth filter and single units were sorted using the Blackrock offline spike sorter or Blackrock online spike sorting software. After sorting, spikes were quantified using Neuroexplorer 5.

### Treatment

Water-soluble CTZ (Nanolight #3031) and CTZ solvent (Nanolight #3031C) were used throughout. For treatment, animals received 30 µL of CTZ (150µg) or equivalent vehicle solvent, including approximately 7-10 µL cannula dead volume. Ventricular infusions were delivered at 4 µL/min every other day for 14 days beginning 1 day post injury. During infusions, animals were allowed to freely move in an open field.

### Behavioral testing

Behavioral testing was done using the Basso, Beattie, and Bresnahan (BBB) rating scale for spinal cord injured rats, where rats are rated on a scale from 0-21, with 0 being completely paralyzed, 10 being the first point where weight bearing steps occur, and 21 having a perfect gait *(31)*. All behavioral testing was done by two blinded observers. If behavior testing occurred on the same day as treatment, behavior testing was done prior to the CTZ mediated stimulation.

### Histology

At 5 weeks post injury, rats were given a lethal dose of Fatal Plus (Vortech Pharmaceuticals), and tissue was collected by transcardial perfusion with cold phosphate buffered saline (PBS) followed by 4% w/v paraformaldehyde solution in PBS. Spinal cords were extracted and incubated in the 4% paraformaldehyde solution at 4°C overnight. Prior to freezing, cords were acclimated to 30% sucrose in PBS w/v for 3 days at 4°C, then flash frozen and stored at −80°C. Thoracic and lumbar regions were embedded in M1 embedding matrix (Fisher Scientific), cryosectioned at 30 µm for thoracic and 50 µm for lumbar regions and mounted directly on positively charged slides for histological staining or fluorescent imaging.

For eriochrome cyanine (EC, Sigma) staining, thoracic sections mounted on slides were air dried, dehydrated, and defatted in graded ethanol solutions (50, 70, 90, 95, 100%, 3 minutes each) followed by xylene (10 minutes), rehydrated in graded ethanol solutions, then incubated in EC solution for 10 minutes*(32)*. Slides were rinsed twice with water and differentiated in 0.5% ammonium hydroxide, then rinsed twice with water. Slides were dehydrated in graded ethanol solutions to xylene and cover slipped with Eukitt mounting media (Sigma). Slides were scanned with a Nikon Coolscan IV slide scanner. Spared white matter was quantified by tracings in ImageJ software by personnel blinded to condition.

### Gene expression

Animals were deeply anesthetized with isoflurane, the spinal cord was dissected out, rinsed in cold PBS, and the lumbar enlargement placed into a tube and flash frozen in liquid nitrogen. Samples were stored at −80°C until processed. RNA was extracted using an All Prep kit (Qiagen) per the manufacturer’s instructions. Quantitative PCR (qPCR) was performed as previously described *(33)*. Briefly, complementary DNA synthesis was performed using the High Capacity RNA-cDNA kit (Applied Biosystems). All samples were analyzed in triplicates using a StepOnePlus Real-Time PCR machine (Applied Biosystems) using Eva Green PCR Master Mix (MidSci) in a total volume of 20 µL. Gene expression was normalized to Glyceraldehyde 3-phosphate dehydrogenase (GAPDH). Results were analyzed using the double delta CT method and are expressed as fold expression of sham animals.

### Statistics

All statistical tests (except for RNAseq) were performed in SPSS Statistics 24 (IBM). A two way repeated measures ANOVA was used for BBB with Bonferonni post hoc test *(31)*. For all other analysis, a one way ANOVA with Bonferonni post hoc was used. Sample sizes were estimated using power analysis with G*power 3.1*(34)*.

### RNAseq

Animals were injured, treated, tissue collected and RNA isolated as described above. RNA-Seq Libraries were prepared and sequenced using 75bp single reads in the next generation sequencing core at SCRIPPS Research Institute using the Illumina platform (San Diego, CA). (elaborated in supplemental methods)

## List of Supplementary Materials

### Methods

Figure S1. Cannula design.

Figure S2. Return of bladder function.

Figure S3. Locomotor recovery with luciferase only vs vehicle. Figure S4. Scatter plot for all genes in thoracic region.

Figure S5. Scatter plot for all genes in the lumbar region.

Figure S6. Heatmaps for thoracic and lumbar up and down regulated genes for treatment vs vehicle groups.

Table S1. List of primers used.

Table S2. Pearson’s correlation coefficient for all genes in the thoracic region.

Table S3. Pearson’s correlation coefficient for all genes in the lumbar region.

## Supplementary files

Excel file RNAseq.

## Funding

This study was supported by the National Institutes of Health (U01NS099709), the National Science Foundation (DBI-1707352), the W.M. Keck Foundation, and the CMU Office of Research and Graduate Studies. MP is a W. M. Keck Foundation Fellow.

## Author contributions

EDP and UH designed the experiments and wrote the paper. EDP initiated and conceptualized the study and carried out all experiments and analysis. EDS performed animal care and treatment. AP performed animal care and surgery. LOS performed animal care and treatment. JRZ performed animal care, behavior and treatment. AJP performed histology and analysis. AA performed RNAseq data analysis. MP performed behavior. UH supervised all aspects of the work.

## Competing interests

The authors declare no competing financial interests.

## Data and material availability

All plasmids used in this study will be available from Addgene (pAAV-hSyn-LMO3: 114099; pAAV-Hb9-LMO3: 114103; pAAV-hSyn-sbGLuc-B7-EYFP: 114110). RNAseq dataset is available in Supplementary Material. All other data is freely available for research upon request.

