## Supplemental Material for "Restoring Function After Severe Spinal Cord Injury Through Bioluminescence-Driven Optogenetics"

**List of Supplementary Materials**

Methods
Figure S1. Cannula design.
Figure S2. Return of bladder function.
Figure S3. Locomotor recovery with luciferase only vs vehicle.
Figure S4. Scatter plot for all genes in thoracic region.
Figure S5. Scatter plot for all genes in the lumbar region.
Figure S6. Heatmaps for thoracic and lumbar up and down regulated genes for treatment vs vehicle groups.
Table S1. List of primers used.
Table S2. Pearson's correlation coefficient for all genes in the thoracic region.
Table S3. Pearson's correlation coefficient for all genes in the lumbar region.

**Supplementary files.**

Excel file RNAseq.

**Supplementary Methods.**

RNAseq

RNA-Seq data was tested in Matlab for differentially expressed genes using a negative binomial model*(35)*. A typical differential expression analysis of RNA-Seq data consists of normalizing the raw counts and performing statistical tests to reject or accept the null hypothesis that two groups of samples show no significant difference in gene expression. Here, we used the negative binomial model to achieve that. The starting point for this analysis of RNA-Seq data is a count matrix, where the rows correspond to genomic features of interest, the columns correspond to the given samples and the values represent the number of reads mapped to each feature in a given sample. Here, we used the TPM counts for genes from both the treated and vehicles and lumbar and thoracic spinal cord regions*(36)*.

In order to account for multiple testing, we performed a correction (or adjustment the Benjamini-Hochberg adjustment) of the p-values so that the probability of observing at least one significant result due to chance remains below the desired significance level. We set a threshold of 0.05 for the adjusted P-values, equivalent to considering a 5% false positives as acceptable, and identify the genes that are significantly expressed by considering all the genes with adjusted p-values below this threshold*(37)*.

We identified the most up-regulated or down-regulated genes by considering an absolute fold change above a chosen cutoff. For example, a cutoff of 1 in log2 scale was used which yields the list of genes that are up-regulated with a 2 fold change.

**Supplementary Figures.**

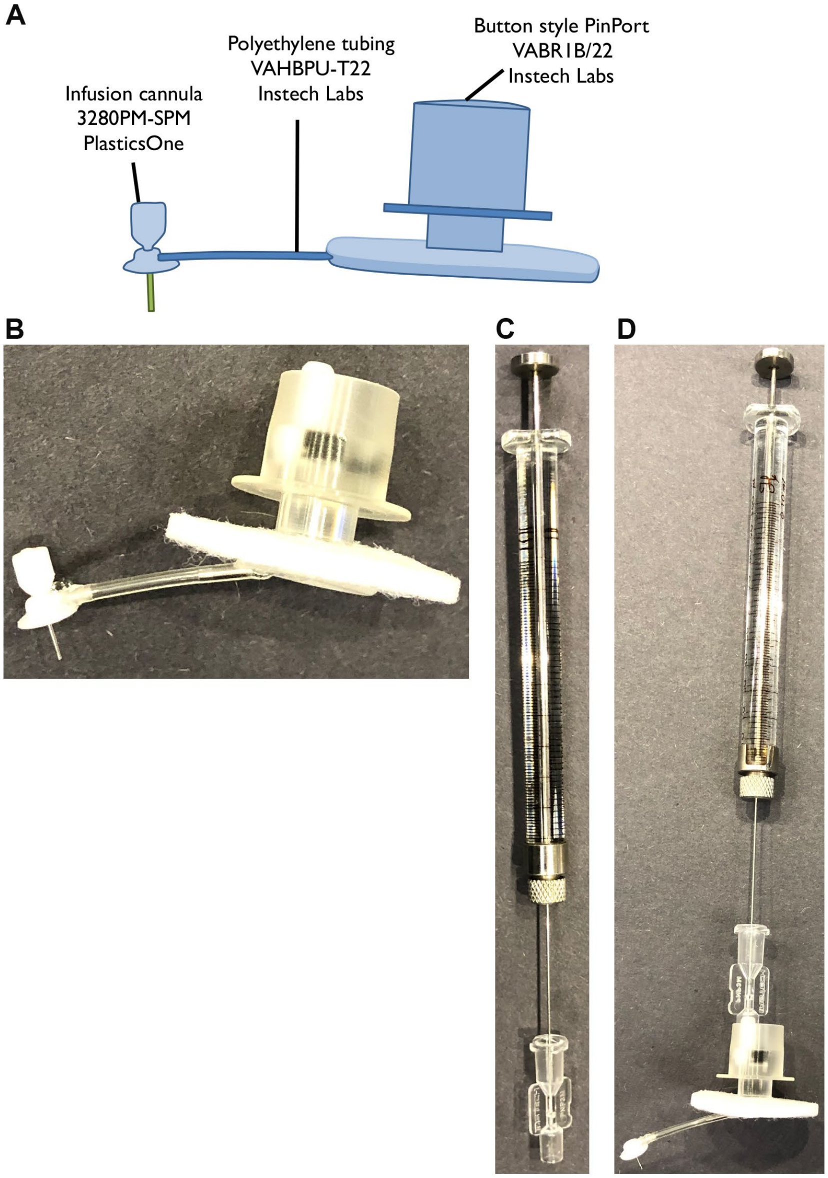


**Fig. S1. Rat lateral ventricle cannula.** (**A**) Schematic of the cannula used in this study consisting of an infusion cannula and PinPort button which are attached by polyethylene tubing and glued together. (**B**) Picture of the assembled cannula. (**C**) A 100µL Hamilton Gastight syringe with a modified 25G blunt needle. To assemble the needle, the metal insert of an Instech PinPort injector was removed, the hole in the center was expanded all the way through with a 25G beveled syringe needle, and the 25G blunt Hamilton needle is inserted to replace the original metal piece with the knurl positioned on the needle first. (**D**) The assembled cannula with injector inserted as it would be for an infusion with this cannula. This approach allows researchers to acutely infuse substances into the brain without restraining the animal as would be necessary with other cannulation approaches.

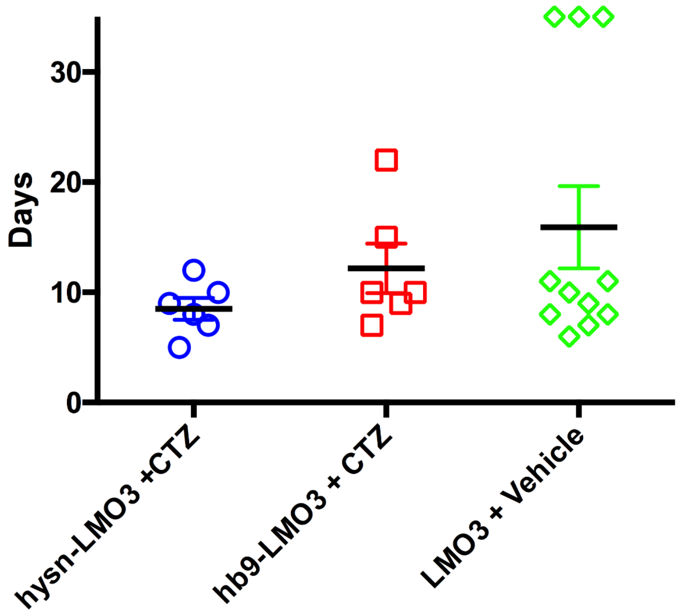


**Fig. S2. Improved bladder function with BL-OG.** Number of days until the return of bladder function are plotted. The groups which received vehicle treatment did not regain bladder control as soon as those receiving CTZ, however this difference was not considered significantly different.

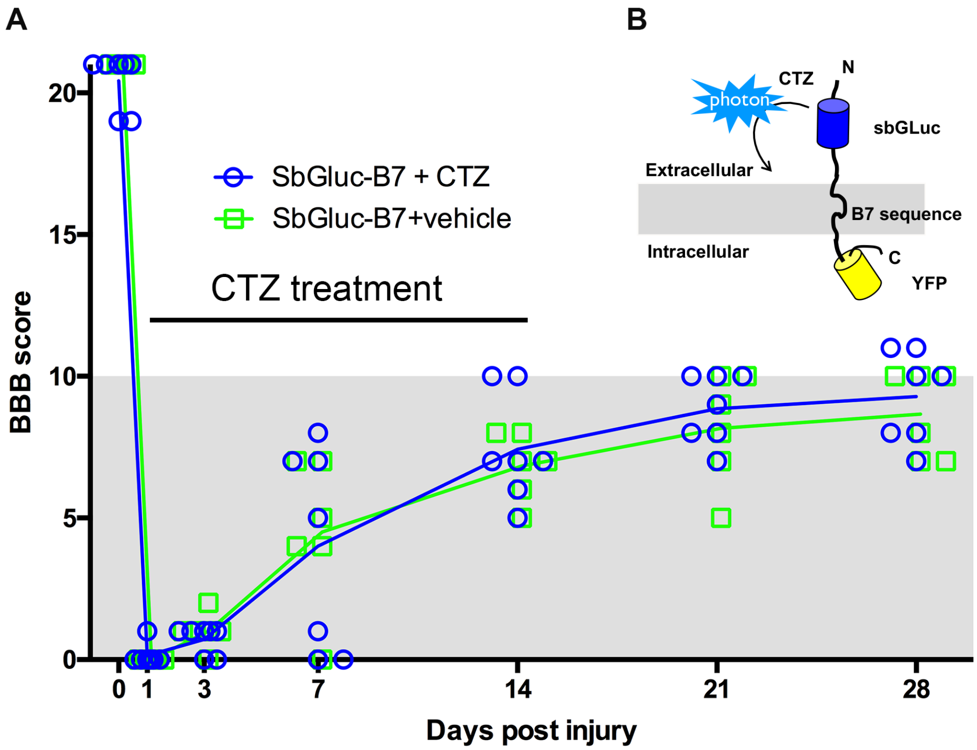


**Fig. S3. Bioluminescence without channelrhodopsin has no effect on locomotor recovery.** (**A**) BBB locomotor scores for rats expressing AAV 2/9 hSyn-sbGluc-B7-EYFP and given CTZ or vehicle following injury. Unlike those treated with CTZ and expressing the opsin, rats only expressing the luciferase component that received CTZ did not recover to a greater extent than the vehicle treated group. (**B**) Illustration of the luciferase only control construct used where the luciferase is extracellular and will produce bioluminescence just as the LMO constructs but will not cause a light sensitive channel to open.

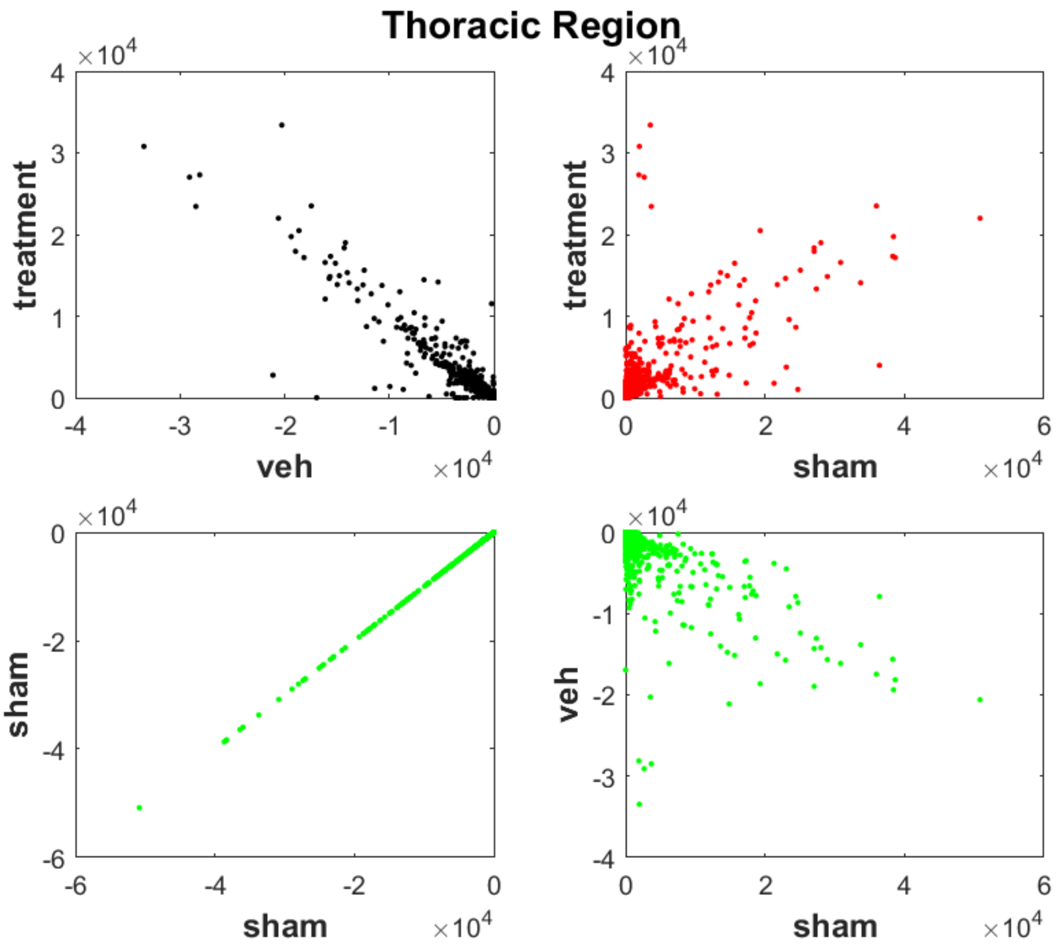


**Fig. S4. Altered gene expression in the thoracic region is a result of injury.** Scatter plots for different pairs of subjects for all genes in the thoracic region. This shows that the treatment and vehicle groups are the most correlated in this region while the sham and vehicle are the least (sham-sham is shown for comparison to a case where correlation coefficient is 1). This indicates that in the thoracic region, the gene expression is dominated by genes that have altered expression as a result of injury to the spinal cord.

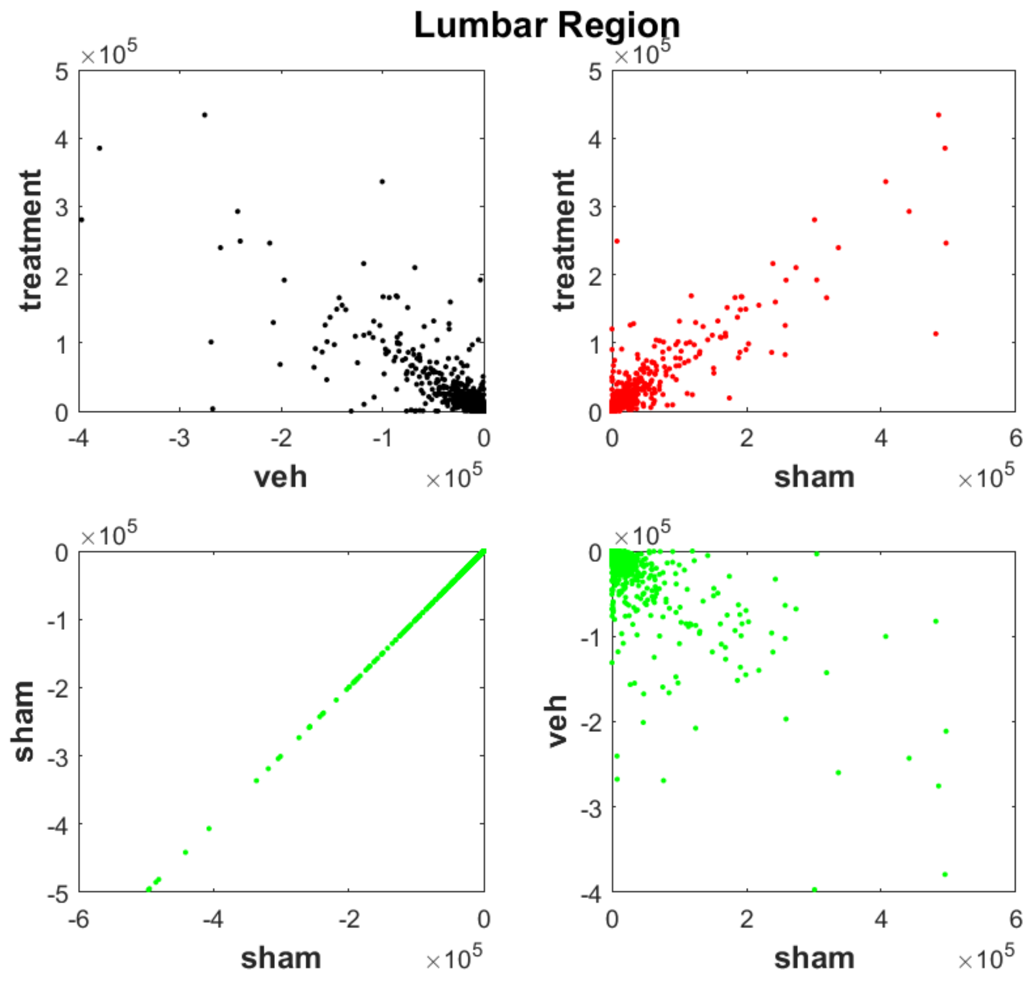


**Fig. S5. Altered gene expression in the lumbar region is a result of neural stimulation.** Scatter plots for different pairs of subjects for all genes in the Lumbar region. Here, the highest correlation is between the treatment and sham group and least between the sham and vehicle group (sham-sham correlation is shown only to highlight what a perfect correlation of 1 would look like). This indicates that stimulation following injury alters gene expression profiles in the lumbar region.

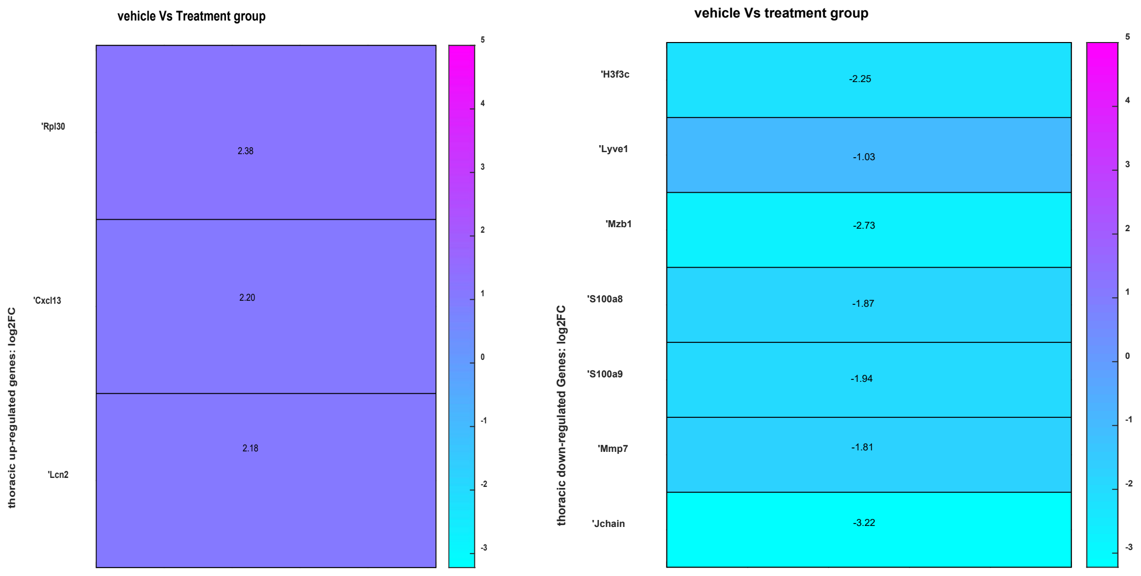


**Fig. S6. Relative expression of selected genes at injury site.** Heat maps for differential gene expression in the thoracic region between treatment and vehicle groups, log2FC values are shown as numbers on each graph.

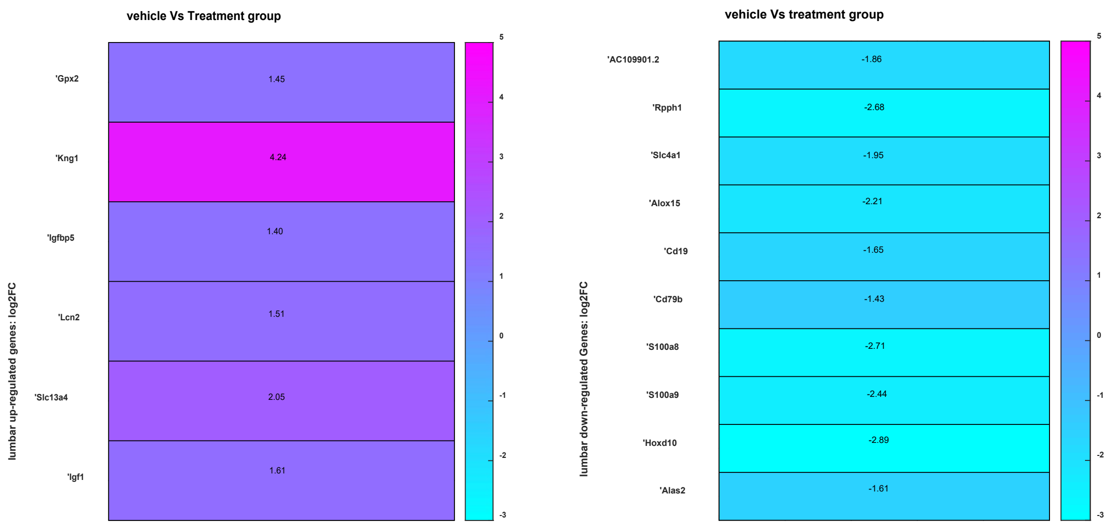


**Fig. S7. Relative expression of selected genes at neural stimulation site.** Heat maps for differential gene expression in the lumbar region between treatment and vehicle groups

**Supplementary Tables.**

**Table S1. List of primers used.**

| Gene | Forward 5'-3' | Reverse 5'-3' |
|---|---|---|
| GAP-43 | CAAGCTGAGGAGGAGAAAGAAGC | GGAGAGACAGGGTTCAGGTG |
| Map2 | CACTGGAAGAAGCCTCGAAGATG | TCCTTGTCTAAAGGCTCAGCG |
| NMDAR2d | ACCCTGACATGCACAGCTAC | TCCAGTTTCCCTGCCTTGAG |
| PSD-95 | AACACGGACACCCTAGAAGC | CAGACCTGAGTTACCCCTTTCC |
| BDNF | GAAGAGCTGCTGGATGAGGAC | TTCAGTTGGCCTTTTGATACC |
| VEGF | CAGAAAGCCCATGAAGTGGTG | GGGCTTCATCATTGCAGCAG |
| Caspace-3 | GAGCTTGGAACGCGAAGAAAAG | AGAGTCCATCGACTTGCTTCC |
| Bcl-2 | GGATAACGGAGGCTGGGATG | AGCAGCGTCTTCAGAGACAG |
| iNOS | ATCTTGGAGCGAGTTGTGGATTGTT | GGTAGTGATGTCCAGGAAGTAGGTGA |
| Arginase | TGTGGTAGCAGAGACCCAGAAGAAT | CAGCGGAGTGTTGATGTCAGTGT |

**Table S2. Pearson's correlation coefficient (r ) for all genes in the thoracic region.**

| **Thoracic region** | **r** |
|---|---|
| Treatment-vehicle | 0.94 |
| Sham-vehicle | 0.73 |
| Sham- treatment | 0.75 |

The treatment and vehicle group show the highest correlation while there is little difference between the sham-vehicle and sham-treatment groups. This indicates that the similarity could be due to injury to this region of the cord which was inflicted on the two groups other than the sham.

**Table S3. Pearson's correlation coefficient (r ) for all genes in the lumbar region.**

| **Lumbar region** | **r** |
|---|---|
| Treatment-vehicle | 0.88 |
| Sham-vehicle | 0.79 |
| Sham-treatment | 0.90 |

The sham-treatment group show the highest correlation while the sham-vehicle group show the least. This could be because the gene expression in treatment group was following the sham group while that of vehicle group was the most dissimilar which could indicate it was related to higher expression of genes producing healthy proteins.
